# Whole genome sequence and 16S rRNA gene amplicon metagenomics of enhanced in-situ reductive dechlorination at a tetrachloroethene-contaminated superfund site

**DOI:** 10.1101/2024.06.20.599892

**Authors:** Rebecca A. Reiss, Peter A. Guerra, Oleg Makhnin, Matthew Kellom

## Abstract

The application of environmental DNA analysis techniques to guide the bioremediation strategy for tetrachloroethene-contaminated groundwater is exemplified by the North Railroad Avenue Plume (NRAP) Superfund site located in New Mexico, USA. Enhanced reductive dechlorination (ERD) was selected as the remedy due to the presence of tetrachloroethene biodegradation byproducts, organohalide respiring genera *Dehalococcoides* and *Dehalobacter*, and associated reductive dehalogenase genes detected prior to remediation. DNA extracted from groundwater samples collected prior to remedy application and after four, 23 and 39 months was subjected to 16SrRNA gene amplicon and whole genome sequencing (WGS). The goals were to compare the potential of these methods as tools for environmental engineers and to highlight how advancements in DNA techniques can be used to understand ERD. The response of the indigenous NRAP microbiome to the injection and recirculation of electron donors and hydrogen sources is consistent with results obtained from microcosms, dechlorinating consortia, and other contaminated sites. WGS detects three times as many phyla and six times as many genera as 16S rRNA gene amplicons. Both techniques reveal abundance changes in *Dehalococcoides* and *Dehalobacter* that reflect organohalide form and availability. No methane was detected before remediation, its appearance after biostimulation corresponds to the increase in methanogenic *Archaea*. Assembly of WGS reads produced scaffolds containing reductive dehalogenase genes from *Dehalococcoides*, *Dehalobacter, Dehalogenimonas, Desulfocarbo,* and *Desulfobacula*. Anaerobic and aerobic cometabolic organohalide degrading microbes that increase in abundance at NRAP include methanogenic *Archaea*, methanotrophs, *Dechloromonas*, and *Xanthobacter*, some of which contain hydrolytic dehalogenase genes. Aerobic cometabolism may be supported by oxygen gradients existing at the aquifer-soil interface or by microbes that have the potential to produce O_2_ via chlorite dismutation. Results from next-generation sequencing-based methods are consistent with current hypotheses regarding syntrophy in environmental microbiomes and reveals novel taxa and genes that may contribute to ERD.

## Introduction

### Tetrachloroethene prevalence and microbial biodegradation

The use of tetrachloroethene (PCE) for dry cleaning began in the 1930s but health concerns were not recognized until the 1960s [1]. Contamination of groundwater with PCE from its use in dry-cleaning and in degreasing threatens potable water supplies worldwide [2]. The complete microbial anaerobic respiration of PCE consists of a sequential process in which successive reductase enzymes remove chlorine atoms (Cl) to ultimately yield ethene, which is readily mineralized by soil microbes and is not known to be a carcinogen. Chlorinated volatile organic compounds (cVOCs) such as PCE and its degradation byproducts vary in toxicity [3]. PCE is a probable carcinogen (group 2A) for all routes of exposure: inhalation, ingestion, or dermal. PCE reductase enzymes cleave the first Cl, generating trichloroethene (TCE), a carcinogen (group 1) regardless of exposure route. Dichloroethene (DCE, cis and/or trans isomers), vinyl chloride (VC), and ethene are formed in succession. DCE carcinogenicity remains undetermined, however, exposure can cause central nervous system depression, liver damage, and infertility. VC is carcinogenic (group 1) by any route of exposure. During each step in the reduction byproduct solubilities increase and their adsorption to soil decrease, increasing their mobility and concentration in groundwater. Complete reduction to ethene without stalling is a key aspect of successful enhanced reductive dechlorination (ERD), otherwise bioremediation could result in an increase in exposure to toxic byproducts.

Organohalides (OHs), including VC, are produced during natural biogeochemical processes and microbial transformations of chlorinated solvents contribute to the global Cl cycle [4, 5, 6]. Dehalogenation mechanisms are widespread in nature and account for the detection of taxa capable of respiring OH at sites undergoing natural attenuation [7, 8]. Enhanced in-situ reductive dechlorination (ERD) involves adjusting groundwater to anaerobic reduced conditions, which encourages microbes containing reductive dehalogenase (*rdh)* genes to respire chlorinated solvents. There are currently 30 bacterial genera validated in the literature that include strains capable of dechlorinating ethenes through anaerobic respiration (Table S1). Collectively, *rdh* gene products regardless of the substrate used, make up the Protein family (Pfam) 13486, based on the alignment of amino acids common to proteins with RD function. Syntrophy involving OH-respiring genera *Dehalococcoides* (*Dhc*) and *Dehalobacter* (*Dhb)* is well-established, *Dhb* dominates during the presence of PCE, TCE, and DCE (also chlorinated ethanes), *Dhc* increases as VC becomes available [9].

The role of *Geobacter* in ethene biodegradation is complex. This genus is known for the transfer of electrons to other microbes along specialized pili [10]. *Geobacter lovleyi* and *G. thiogenes* are distinguished by the presence of *rdh* genes in these genomes, and both are assigned to the genus *Trichlorobacter* in Joint Genome Institute’s (JGI) Integrated Microbial Genome/Microbiomes (IMG/M) database. This is based on the legitimate and validly published proposal to establish a new genus [11]. *Geobacter* can produce cobalamin that can be transferred to *Dhc*, a cobalamin auxotroph [12]. *Sulfurospirillum*, *Acetobacterium* and *Desulfitobacterium* are also cobalamin producers found in dechlorinating consortia [13, 14], exemplifying the importance of syntrophy in the utilization of OH as an energy source [15].

Dechlorination occurs through pathways other than respiration, referred to as cometabolism [16]. The list of genera capable of cometabolizing OH (Table S2) includes methanogenic *Archaea,* first observed in association with dechlorinating communities in the 1980s [17]. In the 1990s it was reported that methanogens harbor reduced transition-metal cofactors capable of dechlorinating TCE in anaerobic conditions [18, 19]. Meta-analysis of OH contaminated sites confirms the linkage between reductive dechlorination and methanogenic taxa [20]. Once PCE is reduced to TCE, it can be a substrate for oxidation by methane and toluene monooxygenases (*mmo* and *tmo*, collectively *mo*), harbored by methanotrophs, forming unstable epoxides [21, 22, 23]. Although the anaerobic conditions for ERD should exclude aerobic cometabolism, oxygen gradients may exist in microenvironments where the aquifer is in contact with the ground [16]. Another possibility is the presence of microbes capable of producing O_2_ in the dark through chlorite dismutation [24]. Either way, epoxide detection is difficult and the fate of these molecules is unclear. If converted into a chlorinated ethanes, these can be respired by strains of *Dhb*, *Trichlorobacter*, and *Desulfitobacterium* [25, 26, 27]. Hydrolytic dehalogenase genes (*hdh*) are diverse, widespread in nature, and harbor the potential to complete dechlorination [7].

### History of the North Railroad Avenue Plume (NRAP)

NRAP was the result of leakage of PCE from a dry-cleaning facility into the sole-source potable water aquifer for the city of Española New Mexico and the Santa Clara Pueblo. TCE and DCE commingled with petroleum hydrocarbon contamination in the plume, which was declared an Environmental Protection Agency (EPA) Superfund Site in 1999 (National Priority List #NMD986670156). The water table is encountered at approximately 1.5 m below ground surface (bgs) in the shallow aquifer, which is comprised of two layers, a high-permeability sand/gravel/cobble mixture, which extends to approximately 6.1 m bgs and a 1.2- to two-meter-thick sequence of interbedded fine-grained sands and sandy clay layers from approximately 6.1 to eight m bgs. Beneath the shallow aquifer is a six- to 12-meter-thick clay layer that is generally continuous across the site and separates the shallow aquifer from the deep aquifer. The lithology below of the shallow aquifer consists of sequences of silts and clays interbedded with fine-grained silty sand and sand units. Additional details of the site geology are available [28].

A pilot test conducted in 2006 determined the optimal amendments and nutrients and their delivery methods for successful remediation [29]. Daughter of PCE degradation were detected and the results of quantitative polymerase chain reaction (qPCR) revealed the presence of *Dhc*, *Dhb,* and *rdh* genes. This data supported biostimulation to achieve ERD, which is achieved by the injection and recirculation of electron donors and nutrients in-situ to deoxygenate and reduce the aquifer in support of anaerobic biodegradation [30]. Bioaugmentation, where non-native microbes are injected, was deemed unnecessary. During the pilot test phase, emulsified vegetable oil (EVO) and hydrogen gas (H^+^) were injected into the source area and dairy whey was injected into the hot spot, which had approximately 10-fold lower concentration of OH. Due to its superior performance, EVO was injected into both areas at the start of full-scale operations.

Prior to remediation total organic carbon (TOC) was low and PCE was the dominant cVOC. TOC increased as a results of EVO injection and after 16 months all daughter products of PCE degradation were apparent. PCE was not detectable at 38 months and ethene was the predominate VOC [29]. As of 2020, 90% of the PCE at the source area was eliminated and concentrations in the rest of the shallow aquifer, including the hotspot, were below Federal Maximum Contaminant Levels [28]. The deepest zones still contain significant amounts of contamination. NRAP is half-way through the expected 30-year completion of OH remediation.

### Application of metagenomics to environmental microbiomes

Whole-genome sequencing (WGS, a.k.a. shotgun metagenomics) involves the random fragmentation of DNA, which is subsequently size selected, primer-adapters are added to both ends, and sequenced then using Next-Generation Sequencing (NGS) instruments. Primer adapters are DNA sequences that provide common markers for DNA sequencing reactions ensuring that subsequent amplification steps are not biased by microbial sequences. Primer adapters also include bar codes that facilitate post-sequencing sample identification so samples can be pooled for automated sequencing. After sequencing, reads are assigned a taxonomic identity, and can be assembled into scaffolds that are long enough to annotate complete functional genes. If adequate coverage is available, further assembly can yield complete genomes, known as metagenome-associated genomes (MAGs) that become new reference genomes irrespective of the availability of cultured representatives.

The application of single-cell sequencing also contributes to the increase in reference genomes to which new sequencing data can be compared. These techniques detect previously unknown diversity, known as the ‘microbial dark matter’ that remains a challenge for phylogenetic annotation [31]. Current standards required a cultured microbe that represents the taxa to be considered ‘validly published.’ Taxa designated as ‘*Candidatus*’ include uncultured reference microbes from MAGs or single-cell reference genomes. Strain designations are initially assigned to microbes in culture, pure or mixed, and can be associated with a function, such as OH respiration. Once the DNA from a strain is sequenced, assembled, annotated, and passes quality control measures, it becomes a reference genome that is used by algorithms to determine the phylogenetic lineage of new sequence data. Many reference genomes do not have validated functional data in the literature, but function can be inferred from the lineage and/or presence of key genes, such as *rdh.* Once sequenced, the genome name is the same as the strain designation. WGS was used to thoroughly characterize an aquifer in Rifle, Colorado, USA and consisted of 150-bp reads generated by the Illumina platform that significantly expanded our knowledge of microbes in groundwater systems [32]. This project included multiple replicates of different parts of the aquifer at different time points and yielded 47 new phyla, and over 2,500 new genomes, demonstrating the advantage of WGS to investigate groundwater samples. Currently, the primary disadvantage for many environmental projects is the cost of WGS compared with other molecular biological techniques, such as qPCR.

NGS also provides a significant advance for 16S rRNA gene analysis by allowing PCR amplicons to be sequenced *en masse.* Sequences are identified by sequence comparison to 16S rRNA gene databases. Such techniques are subject to bias inherent in PCR since *a priori* knowledge of sequence is required and target molecules compete for the same primers. This method is less expensive than WGS and is a suitable replacement to phospholipid fatty acid analysis (PLFA) and denaturing gradient gel electrophoresis (DGGE) analyses previously used to characterize the functional potential of microbial communities. While 16S rRNA gene amplicons alone cannot confirm the presence of genes important in biodegradation, additional primer combinations specific for genes of interest can reveal their presence. Microarrays to identify *rdh* genes specific to *Dhc* use this strategy [33].

This report covers the first 3.25 years of ERD in the shallow aquifer and includes resequencing with updated techniques from the previously reported baseline and four month samples from the source area [34]. New data include source area samples from 23 and 39 months and hotspot samples from four and 23 months that were subjected to WGS and 16S rRNA gene amplicon analyses (Table 1). Two post-sequencing workflows for WGS short-read data were performed, phylogenetically annotated unassembled reads (WGS-U) and reads assembled into scaffolds (WGS-A) (Fig. 1). Gene annotations for NRAP scaffolds and for all genomes in the IMG/M database were searched to identify key functional genes then merged with the WGS-U timeline data to identify the functional potential of taxa, a process we proposed to call Function-Informed Phylogeny (FIP) to indicate that both literature-validated functional information derived from studies of culturable stains and sequence-derived gene and taxonomic information from genomes that may not be culturable are included.

**Fig. 1.**
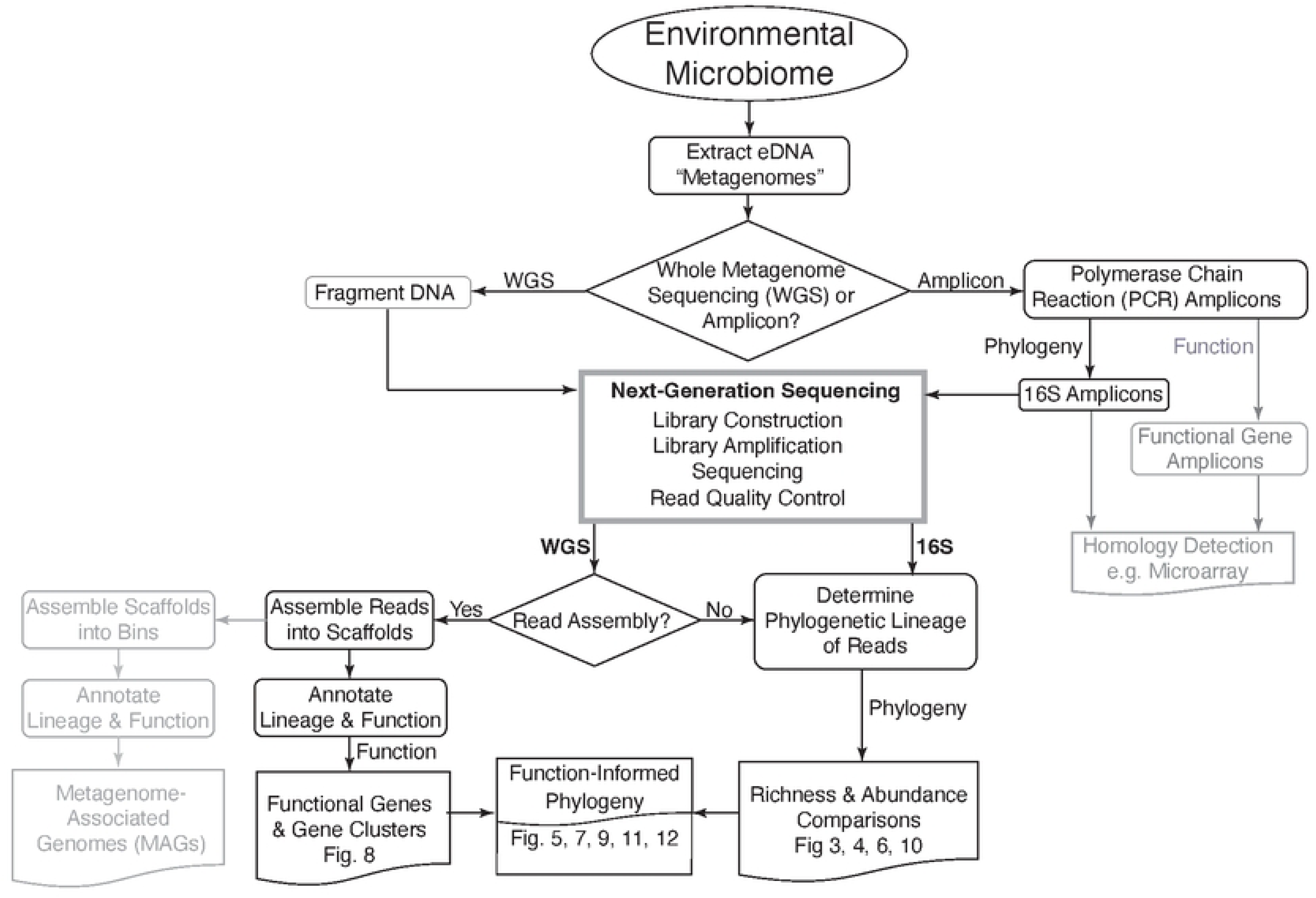
Environmental microbiome metagenomic workflow. WGS and 16S rRNA gene amplicon techniques were applied to the same environmental DNA (eDNA) samples. For WGS, eDNA is fragmented prior to the addition of primer-adapters. For 16S rRNA gene amplicons, eDNA was first subjected to PCR, after which primer-adapter sequences were added to the resulting DNA fragments. NGS protocols were used for WGS and 16S rRNA amplicons. Amplicon sequences were compared to a 16S rRNA gene database for taxonomic identification. For WGS, comparison to the IMG/M database determined lineage. Function-informed phylogeny involves combining literature-validated functional information with function inferred by the presence of key genes then mapping the abundance changes to the list, facilitating identification of additional taxa involved in ERD. The steps in grey are not part of this study but are included to indicate other applications and future work.

**Table 1.** NRAP eDNA Shallow Plume Sampling History.

| Well | Samples (Dates) | WGS | 16S |
| --- | --- | --- | --- |
| Source Area Extraction Well 3 (SAE3) | **SAE3-00**: 0 months, Baseline (June 2007) | √ | √ |
|  | **SAE3-04**: 4 months (Nov. 2007)<sup>1</sup> | √ | √ |
|  | **SAE3-23**: 23 months (June 2009) | √ | √ |
|  | **SAE3-39**: 39 months (Oct. 2010)<sup>2</sup> | √ | √ |
| Hotspot Extraction Well 6 (HSE6) | **HSE3-0<sup>3</sup>**: Baseline, 0 months (June 2007) | ND | √ |
|  | **HSE6-04**: 4 months (Nov. 2007)<sup>2</sup> | √ | √ |
|  | **HSE6-39**: 23 months (June 2009) | √ | √ |

ND – Not Done:
1. Listed as SAE3-5 and HSE3-5 in the JGI Integrated Microbial Genomes (IMG/M) database
2. Listed as SAE3-47 in the IMG/M database
3. Insufficient DNA from the HSE6 baseline was isolated for WGS and only one 16S rRNA gene amplicon run was available.

In this report, ‘strains’ refer to microbes in culture, either pure or mixed, that can be assigned a function, such as the ability to respire or cometabolize OH. The term ‘genome’ indicates fully sequenced and annotated strains as listed in IMG/M [35]. Additionally, the use of OH-respiring genera (OHRG) rather than OH respiring bacteria (OHRB) and COMG instead of cometabolic bacteria (COMB) reflects the focus on genus-level taxonomic identification and allows for the inclusion of *Archaea*.

The goals of this project are to compare the efficacy of WGS and 16S rRNA gene amplicon techniques as tools for environmental engineering decisions and to expand the understanding of environmental microbiomes capable of ERD.

## Materials and methods

### Sample collection and DNA preparation

Site sampling (Table 1) involved the previously described on-site filtration of microbes and direct extraction of eDNA in the lab that eliminates enrichment procedures [34]. Biomass estimates were based upon the DNA yield per liter of water filtered. The concentrations of all VOCs and methane for each timepoint was extracted from the EPA site report [29].

### Library preparation, sequencing, and bioinformatics workflow

The eDNA was sequenced and analyzed as part of the JGI Community Science Project 1243 (<u>CSP1243</u>). The JGI Microbial Genome Annotation Pipeline (MGAP) was utilized for the bioinformatics workflow [36, 37].

Standard Illumina TrueSeq^®^ protocols were used for WGS. Once the eDNA was fragmented, the 270 bp fraction was selected, and bar-coded primer-adapters were added to each sample. Only one replicate was available for each sample but includes sequencing from both ends. The samples were pooled and run in a single lane of an Illumina HiSeq-2000 flow cell and paired-end reads of 150 bp were generated using standard operating protocols as recommended by the manufacturer. The MGAP protocol includes quality control, contamination filtering, and read annotation [37]. Unassembled 150 bp reads are translated into six reading frames of 50 amino acids and is assigned a strain-level taxonomic identification (IDed) when at least 30% of the amino acids of a reading frame match a genome from the IMG/M reference database and at least 70% of the gene sequence is covered by the alignment, otherwise the read remains unidentified.

Filtered reads from each of the six WGS metagenomes were assembled individually and were also combined into one assembly and annotated through GMAP. Assembly involves first aligning reads into overlapping contiguous reads (contigs), with or without ambiguous bases. Protein coding regions of scaffolds can then be predicted based on gene sequence models and subsequently annotated according to reference databases of protein sequences. Read mapping to scaffolds produces an overall length and the number of times each base is covered by a read averaged over all bases, the depth. Scaffold size is the length times the depth as measured in bp. Taxonomic assignments are made when the percentage of predicted protein-coding regions in a scaffold exceeds 50% for any taxonomic level. For example, if at least 50% of the coding sequences match the designated species of a reference genome, then the species level assignment is made. However, if species match does not exceed 50%, then the genus-level assignment was made if at least 50% of the coding sequences matched a genus. This approach results in a variety of taxonomic level assignments for scaffolds, from no assignment to the genome-level assignments. This strategy takes advantage of the additional information provided by assembly that is not available in WGS-U, in which a taxonomic assignment is made only if the match to a reference isolate genome is above 70% amino acid identity.

For 16S rRNA gene amplification, eDNA extracts were dispensed into a 96-well plate. Illumina iTAG V4 primers, PCR, and library preparation protocols were followed as recommended by the manufacturer. Sequencing was carried out on the Illumina HiSeq-2000 using the same protocol as WGS. There were 11 sequencing reps for the SAE3 samples, five for HSE6-04, four for HSE6-23, but only one was successful for HSE6-00. The 16SrRNA gene fragments were quality and contamination filtered with FLASH (v1.2.6) and duk (v1.05), respectively. [38, 39] Reads were pair-merged into amplicons with FLASH, then clustered into operational taxonomic units (OTU) with USEARCH (v7.0.959 i86linux32) [40], checked for chimeras with UCHIME [41], and annotated with the Ribosomal Database Project (RDP) Classifier (v2.5) [42].

Raw counts and metadata available through IMG/M and were be downloaded from the <u>JGI</u> <u>Genome Portal</u> and imported into the statistical software package JMP (v17.0.0, Cary, NC) for further analyses. Each line in the downloaded unassembled and 16S rRNA gene read files represents a single read, the data were tabulated at the strain level to determine the number of reads per strain. The files for each well-timepoint were joined. The scaffold data files with the length, depth, and linage information were joined using the scaffold object identifier (OID) as the key. A list of genes coding sequences (CDS) from the combined assembly was downloaded and filtered by searching for keywords (e.g. dehalogenase). The 16S rRNA gene amplicon results are reported at the genus-level with a 95% confidence, so WGS-U was tabulated on the genus for comparison. The WGS-U and 16S rRNA datasets were joined using the genus designation as the matching (key) value. The resulting data files subsequent analyses are provided (File S2).

### Taxonomic Nomenclature Updates

Updating the taxonomic nomenclature was necessary to account for recent changes., including the renaming of 42 phyla. [43] The List of Prokaryotic Names with Standing in Nomenclature (LPSN) was consulted for the most recent assignment. [44] Rapid changes in phylogenetic nomenclature at any level can make the lineages in the databases out-of-date. These steps in JMP were used to update the lineages to ensure that tabulations on any taxonomic level were correct. This process is based on the structure of taxonomic lineages, which is an ontology where any level can have multiple children, but only one parent. For example, all members of the genus *Bacillus* should have the same lineage. Changes in nomenclature at the higher levels can result in different lineages, for *Bacillus* the validly published and correct order name is *Caryophanales*. The validly published synonym, *Bacillales,* is still used in some databases, especially for older entries. The process starts with tabulating the file on the entire lineage and then the resulting file this tabulated on a level (e.g., genus). This produces a list of the number of lineages for each genus, which should be one. Genera with multiple lineages were selected and the lineage corrected based upon the most recent nomenclature. This process of lineage correction, as executed in JMP, is provided in Table S3.

### Normalization to Biomass and Scaling

Once the lineages were corrected, the average DNA yield (µg/L) for each timepoint was used as an estimate of biomass that was used to calculate a normalizing factor (NF) for the sample *i* is found as:

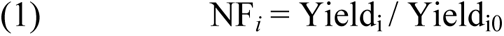

where Yield_i_ = DNA yield (µg/L) for sample *i* and Yield_i0_ is for the lowest-yielding sample (SAE3-00). Read totals by strain (WGS) and by genus (16S rRNA) were divided by this factor to generate normalized reads that were scaled to reads per million (RPM). This value was used for all subsequent tabulations.

### Function-informed phylogeny

Tables S1 and S2 list taxa known to degrade chlorinated ethenes, but metagenomics provides an additional method to identify genomes with this potential by searching for genes that code for the central enzymes involved in dechlorination and other functions. The term function-informed phylogeny (FIP) is proposed to distinguish it from analyses that rely solely on literature-validated comparisons. The filtering and matching processes were done at the reference genome (strain) level. A search of all IMG/M data was conducted starting by entering a term (e.g. ‘dehalogenase’) into the ‘find genes’ function. The resulting files were downloaded and joined with the strain-level WGS-U data. Those that increased in any post-remedy timepoint were considered to have functional potential, regardless of current literature validation. These genome lists were tabulated on the genus level for visualization, excluding strains that lacked the selected function. FIP avoids the limitation of relying solely on genus-level identification of OH-respiring taxa, since some species and strains within a genus harbor *rdh* genes, while others do not. The 16S rRNA gene amplicon data does not make this distinction but the results are shown in the FIP figures for comparison. Results of searches are available (NRAP Genes and Genomes, S4).

### Data Visualization

The standard visualization method used for 16S rRNA gene amplicons is the stacked bar chart with each color representing a different taxon and the abundance represented by the height of the segment. This approach does not work for WGS data since there are too many taxa detected by WGS to make effective use of color to distinguish taxa. Additionally, comparison between samples, including timelines, can be difficult to discern when taxa exhibit large changes in abundance. Cells plots solve these problems since a color gradient is used to indicate abundance, each row is a different taxon, the name is presented next to the row, and timeline data are displayed as columns. Cell plots were generated in JMP after applying binning formulas for each RPM column and are sorted on the basis of abundance at baseline. A dummy variable for each bin was added to ensure that all columns were uniformly scaled, these variables are shown in NRAP Figure Files, but are hidden in the figures. Line graphs of specific genera were generated by selecting the genus in the file containing WGS and 16S rRNA reps, creating a stacked file, and using the Graph Builder function in JMP to overlay the 16S rRNA and WGS-U data for the SAE timepoints. Stacked bar graphs were used for cVOC data and shaded line graph for methane data. The data files used to create cell plots and graphs provided in Excel format (NRAP Figure Files, S3). Image files were exported into Adobe Illustrator (V24.0.1) for markup and file type conversion.

## Results

### Sampling and sequencing overview

At 0 months (baseline), more than 400 liters were filtered without clogging. After remedy application, some filters clogged after 6.25 liters of flow (Fig. 2A). Biomass, as measured by µg DNA per liter exhibited an inverse relationship. *Dhc, Dhb, Desulfuromonas* and *Desulfitobacterium* were detected in the baseline sample with WGS-U, consistent with the original qPCR findings but only 0.2% of baseline reads were IDed, while 2.3% to 5.6% were IDed at all other time points (Figure 2B). Overall, 97% of the reads remained unidentified for all samples. Average scaffold size was greatest at baseline then decreases about 100-fold (Fig. 2C). The number of scaffolds was 100-fold lower in the baseline, compensating for the increased size. There are no baseline scaffolds for *Dhc* or *Dhb* despite their detection by qPCR but both are represented in scaffolds at four, 23, and 39 months. A total of 2.6 x 10^6^ scaffolds resulted from the combined assembly, 0.1% are assigned to a genome, 54% were assigned to a genus, and 35% had no taxonomic ID. Over 4 x 10^6^ CDS were contained within all NRAP scaffolds, representing 30,940 genes, including 161 Archaeal and 1,711 16S rRNA small subunit genes. Only 9.7% of the small subunit genes were assigned to a genus with WGS-A, congruous with the observation that only 3% of all WGS-U reads were assigned to a lineage.

**Fig. 2.**
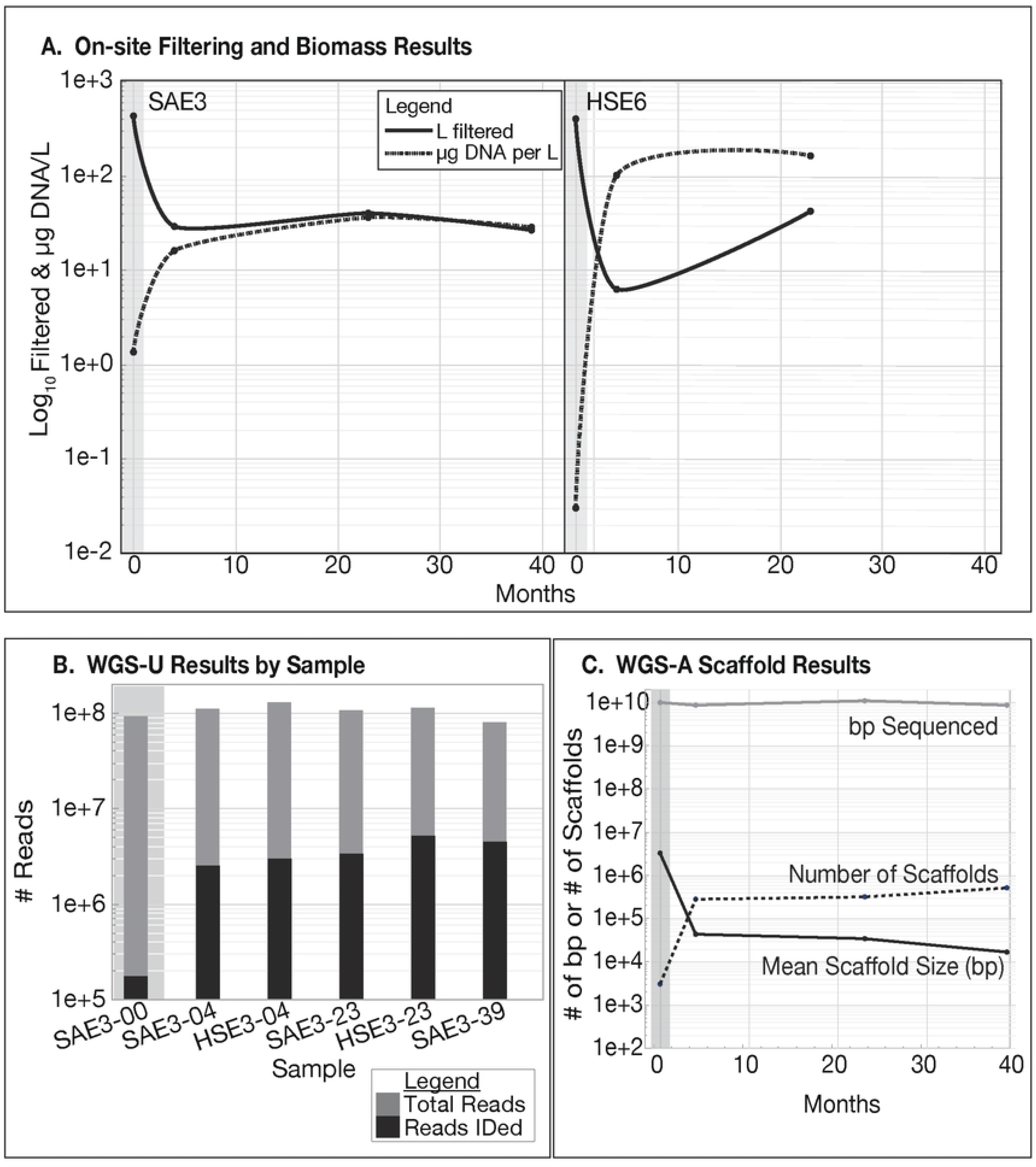
NRAP overview. The baseline (0 months) is shaded and all Y-axes are log_10_ values. **A.** Liters of water filtered on-site and biomass as measured by DNA yield from each sample. **B.** Sequencing depth and WGS-U identification. **C.** Sequencing depth and scaffold profiles for well SAE3 over time.

Amplicons for *Dhc* and *Dhb* were detected in the baseline sample. Taxa richness, as determined by the number of taxa present at each phylogenetic level, was greater in WGS data than 16S rRNA gene amplicon data (Table 2). Both techniques detected higher richness at baseline (SAE3-00) and this observation held for all samples over most taxonomic levels. The one exception is the phylum level 16S rRNA gene amplicon data, where richness was the same at baseline.

**Table 2.** Taxa Richness by Sample and by Method.

|  | Phylum |  |  | Class |  |  | Order |  |  | Family |  |  | Genus |  |  |
| --- | --- | --- | --- | --- | --- | --- | --- | --- | --- | --- | --- | --- | --- | --- | --- |
| Method | 16S<sup>1</sup> | WU<sup>2</sup> | WA<sup>3</sup> | 16S | WU | WA | 16S | WU | WA | 16S | WU | WA | 16S | WU | WA |
| Sample |  |  |  |  |  |  |  |  |  |  |  |  |  |  |  |
| Overall | 19 | 57 | 50 | 38 | 119 | 122 | 69 | 224 | 257 | 127 | 460 | 563 | 241 | 1499 | 2557 |
| SAE3-00 | 18 | 51 |  | 33 | 106 |  | 58 | 205 |  | 98 | 428 |  | 159 | 1358 |  |
| HSE6-00<sup>4</sup> | 15 |  |  | 28 |  |  | 60 |  |  | 106 |  |  | 171 |  |  |
| SAE3-04 | 18 | 44 |  | 30 | 94 |  | 54 | 183 |  | 84 | 373 |  | 136 | 1141 |  |
| HSE6-04 | 17 | 44 |  | 32 | 94 |  | 53 | 183 |  | 83 | 373 |  | 134 | 1142 |  |
| SAE3-23 | 18 | 44 |  | 33 | 94 |  | 58 | 183 |  | 95 | 373 |  | 157 | 1143 |  |
| HSE6-23 | 17 | 44 |  | 35 | 94 |  | 58 | 182 |  | 91 | 372 |  | 145 | 1141 |  |
| SAE3-39 | 18 | 44 |  | 33 | 94 |  | 54 | 183 |  | 103 | 373 |  | 180 | 1143 |  |

1. 16S rRNA gene amplicon
2. WGS-U
3. WGS-A
4. Based on a single replicate of 16S rRNA gene amplification, no WGS data are available.

### ERD from domain to strain

Domain, phylum, class, genus, and strain level analyses reveals the microbiome shifts during ERD. The number of genera attributed to *Archaea* is 113 in WGS-U and nine in 16S rRNA. Fig 3 are domain-level comparison of WGS-U and 16S rRNA gene amplicons compared with methane concentrations.

**Fig. 3.**
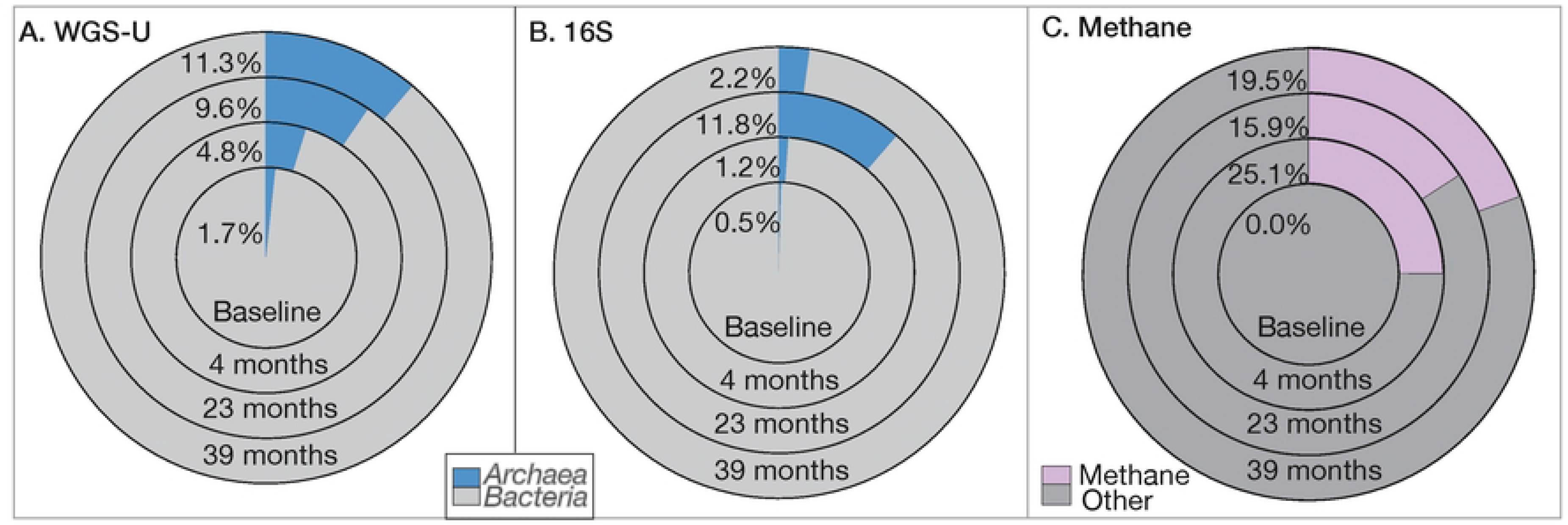
Domain-Level Abundance and Methane Saturation. WGS-U and 16S rRNA gene amplicons detected *Archaea* at baseline but methane was not detected until after the start of remediation.

Overall, 57 phyla were detected by WGS-U, 19 were detected by 16S rRNA gene amplicons (Table 2). *Candidatus* phyla represented 35% of the phyla detected by WGS-U and 0.6% of the reads. Thirteen of the 57 phyla were found only in the baseline (Table S3), representing 1.3% of pre-treatment reads, 12 of these were *Candidatus*. References for *Candidatus* phyla detected at NRAP are provided in Tables S4 and S5. Phyla detectable after baseline are shown in Fig. 4, along with selected class-level abundances.

**Fig. 4.**
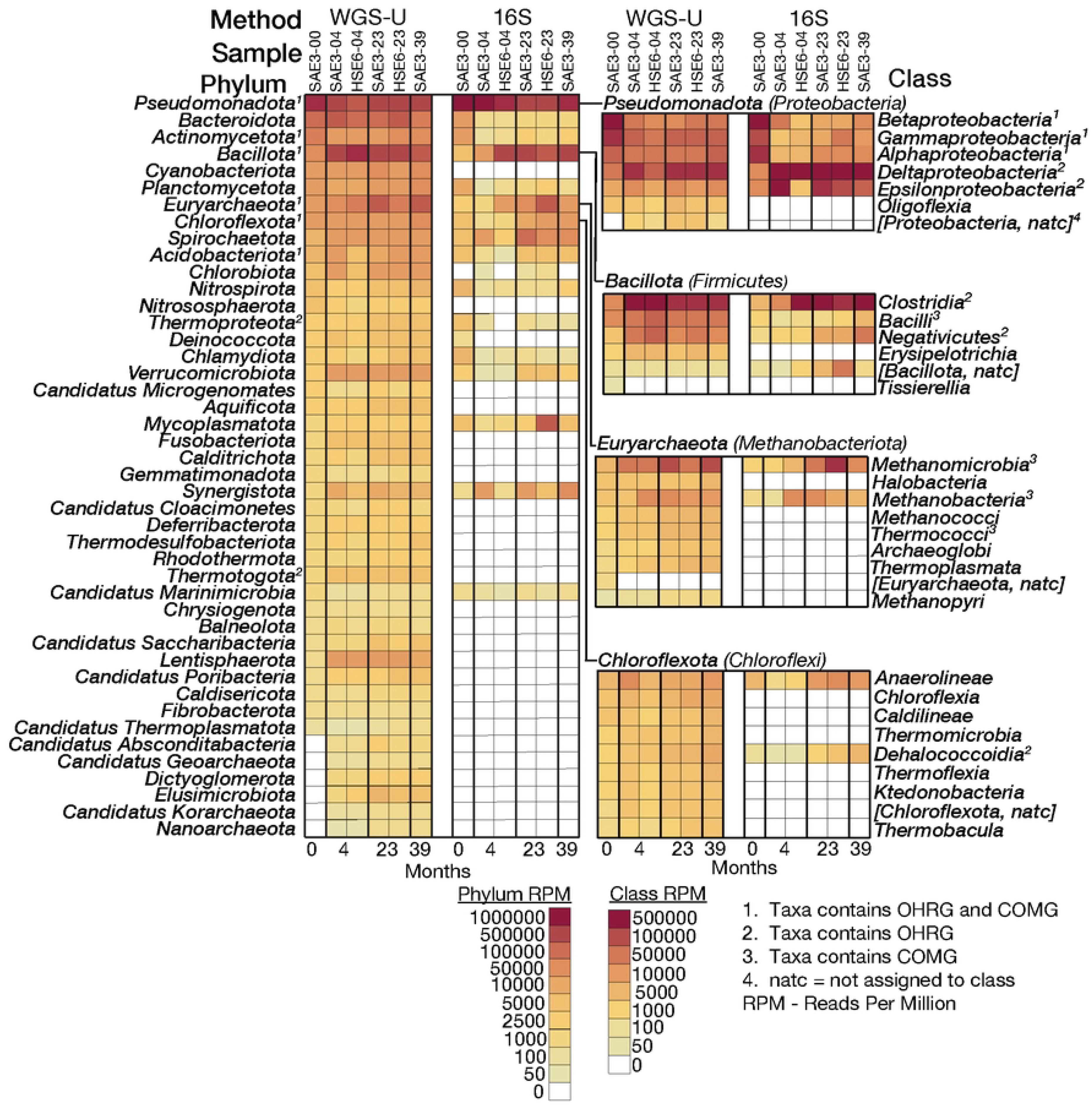
Phylum and Selected Class-Level Abundances. Cell plots are sorted from high to low - abundance at baseline. WGS-U and 16S rRNA data are shown for six samples, including two samples for four- and 23-month sampling events. OHRG and COMG classification include only literature validated taxa (Tables S1 and S2).

There are 2,742 genomes in IMG/M that contain *rdh* genes that encompass 219 genera, only 30 are validated in the literature as OHRGs. Ninety-eight *rdh*-containing genomes are represented at NRAP, 57 increased in abundance overtime and were contained within 32 genera (Fig. 5).

**Fig. 5.**
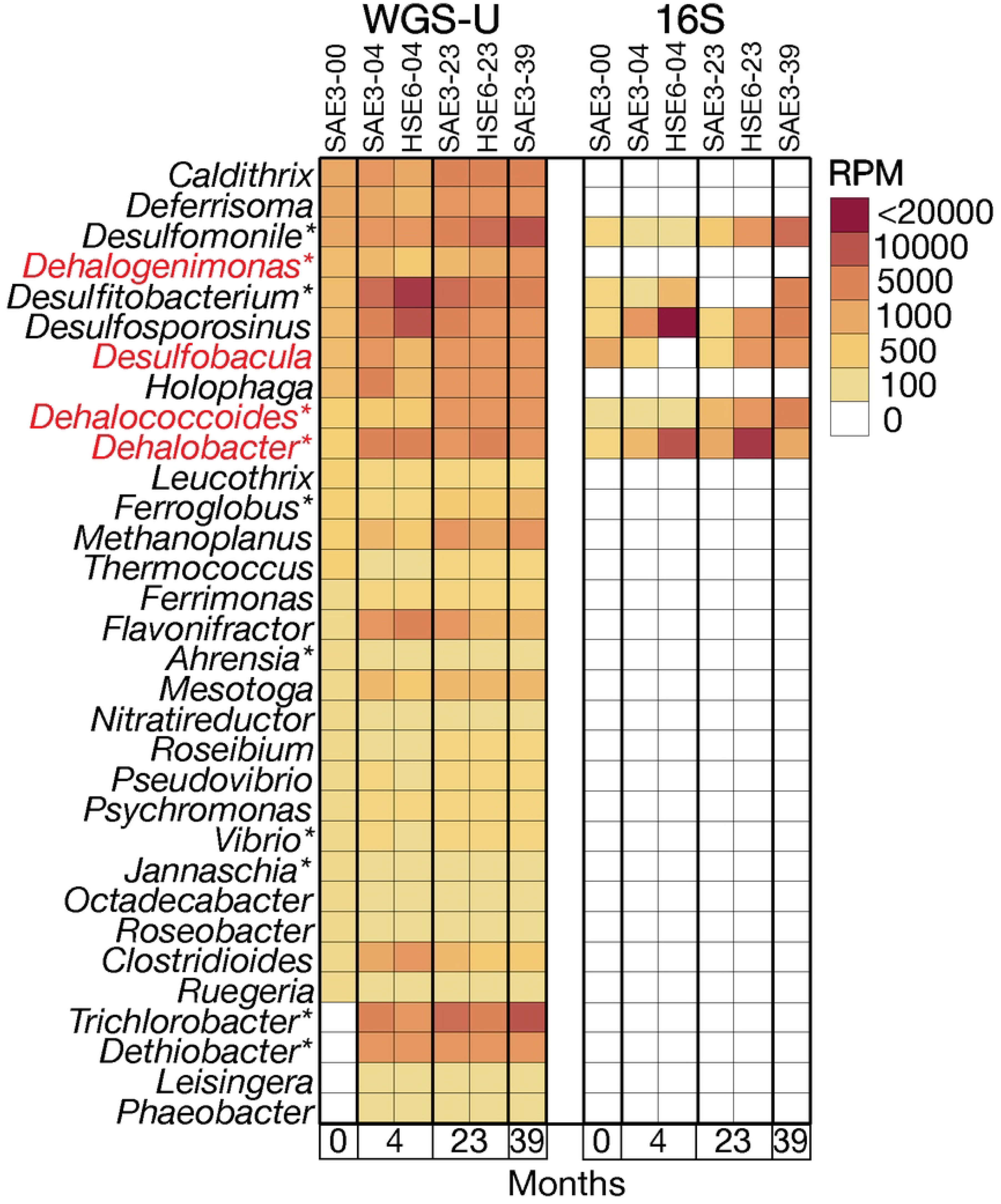
FIP-Selected OHRG at NRAP. These genera increase in abundance at NRAP and have *rdh*-containing genomes in IMG/M. The red font indicates that *rdh* -containing scaffolds were detected at NRAP and asterisks indicate literature-validated OHRG (Table S1).

A comparison of genus-level WGS and 16S rRNA gene amplicon results for selected OHRGs and supporting genera are shown with respect to OH concentrations (Fig. 6). Each WGS sample is represented by one paired-end sequencing run while the 16S rRNA is based on 11 replicates. WGS did not detect *Trichlorobacter* at baseline and was not distinguished by 16S rRNA. *Dehalogenimonas* was not detected at any timepoint by 16S rRNA.

**Fig. 6.**
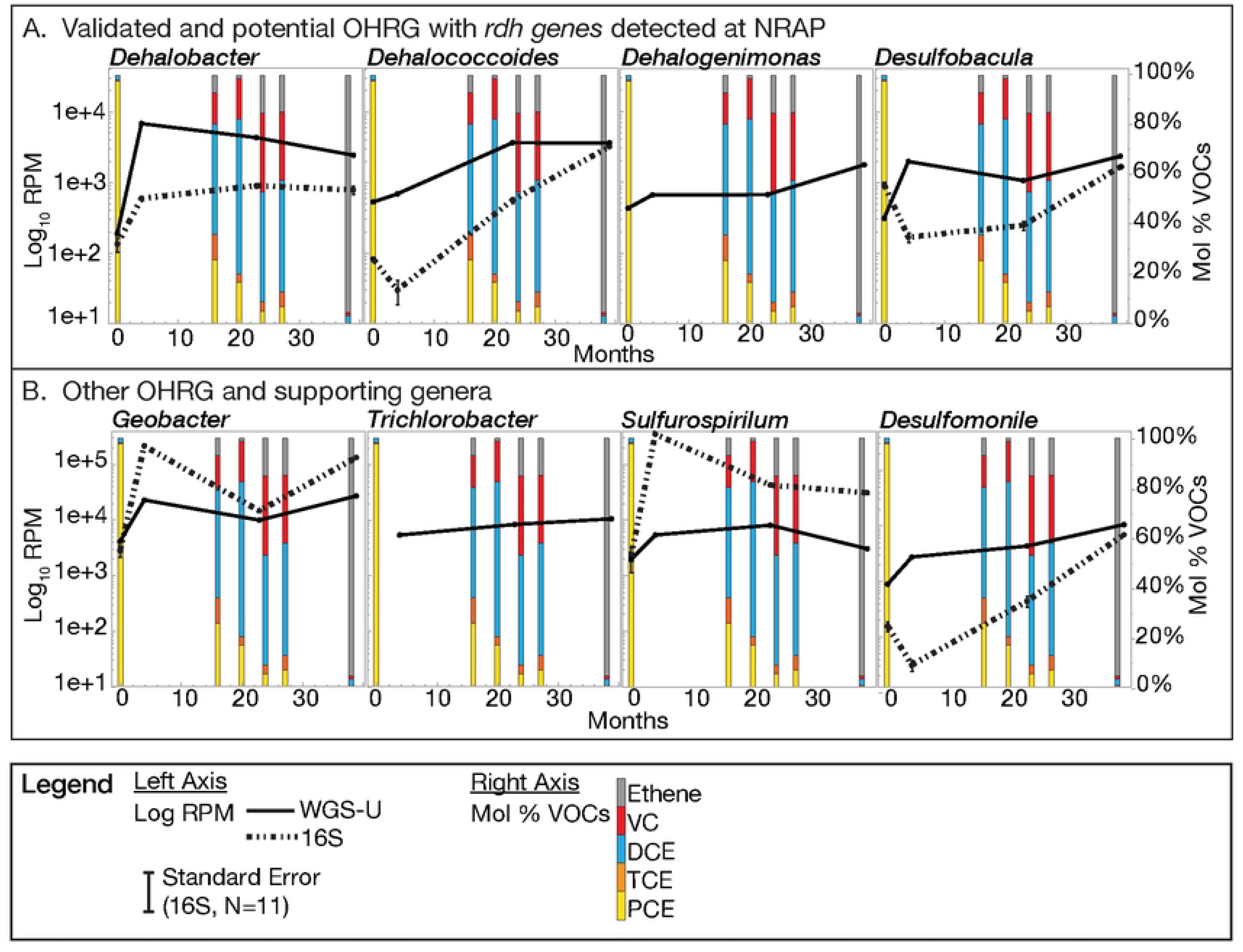
WGS v 16S rRNA: Selected OHRG. The standard errors for the 16S rRNA gene amplicons are shown, but most are too small to be discernable. Note that the RPM range is greater in B than the other graphs.

The profile of chlorinated compound respiration potential (Fig. 7) was based on the functional validation of strains [16] [45]. Combining the OH and abundance profiles demonstrated that PCE and TCE can be respired by 14 strains, DCE by seven, and VC by three, *Dhc. mccartyi 195, BAV1*, and *VS*. The *Sulfurospirillum* species at NRAP lack *rdh* genes and are not known to respire OH. Strains not represented in this list due to the lack of literature validation or were not detected at NRAP.

**Fig. 7.**
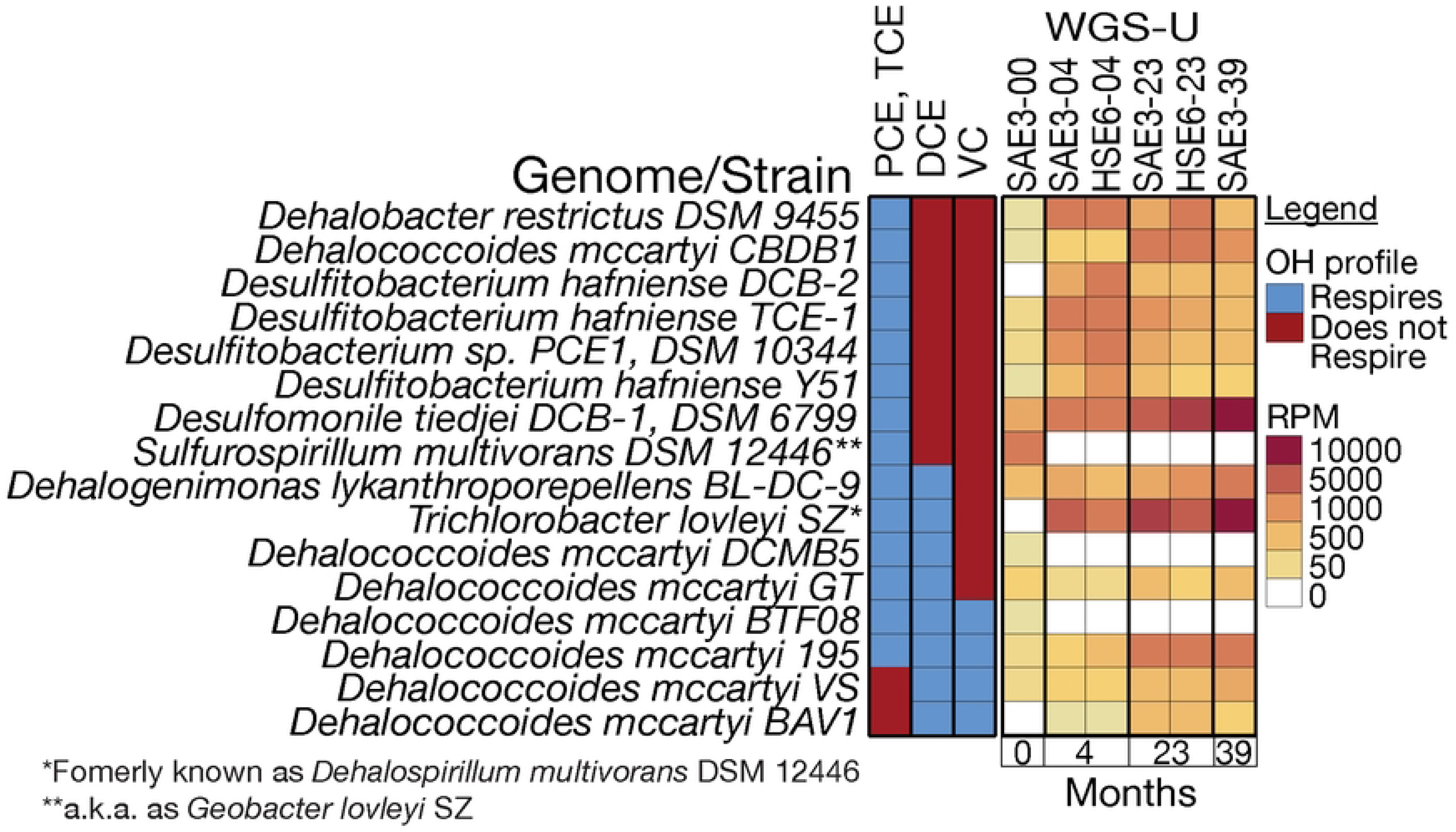
Strain-Level OH Respiration and Abundance Profiles. Literature-validated OH-respiring strains combined with abundance profiles demonstrates the pathway for PCE degradation at NRAP. References for the OH-profiles are provided in Table S6.

The combined assembly provided adequate coverage to detect 390 scaffolds with at least one dehalogenase gene, including 20 *rdh* genes, representing five genera (Table S7). *Dehalobacter*, *Dehalococcoides*, *Dehalogenimonas* and *Desulfobacula* included *rdh*-containing scaffolds at NRAP. *Desulfocarbo* also has a *rdh*-gene containing scaffold but was not detected by WGS-U or 16S rRNA gene amplicons. Scaffolds shown in Fig 8 include *rdh* and genes for accessory proteins necessary for OH-respiration.

**Fig. 8.**
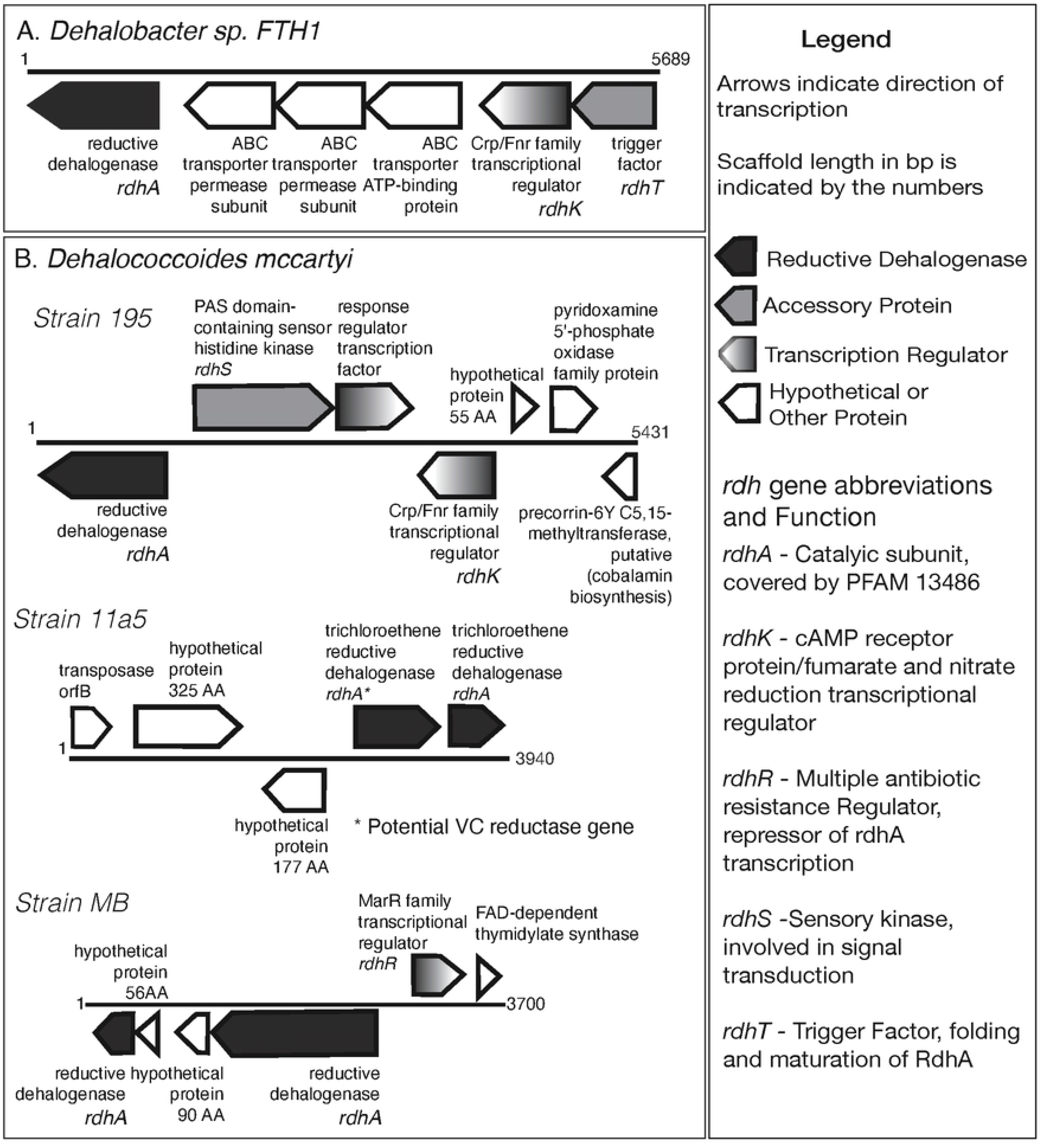
Representative Scaffolds Containing *rdh* Gene Clusters. Scaffolds are the result of a combined assembly that includes all timepoints and both wells. Information for all *rhd*-containing scaffolds is listed in Table S7.

### Anaerobic and aerobic cometabolism at NRAP

The increase in *Archaea* supports anaerobic cometabolism. To determine whether WGS can provide evidence for the involvement of aerobic cometabolic pathways during biodegradation, IMG/M was searched for microbes with key gene products (e.g. monooxygenases) and the phylogenetic data joined with NRAP abundance data. There are 650 instances of *tmo* and *mmo* genes listed in IMG/M in 539 genomes, 78 of these are found at NRAP and represent 40 genera. Not all of these microbes are classified as methanotrophs but harbor the potential to oxidize TCE. The 19 shown in Fig. 9 increase in abundance over baseline in at least one sample.

**Fig. 9.**
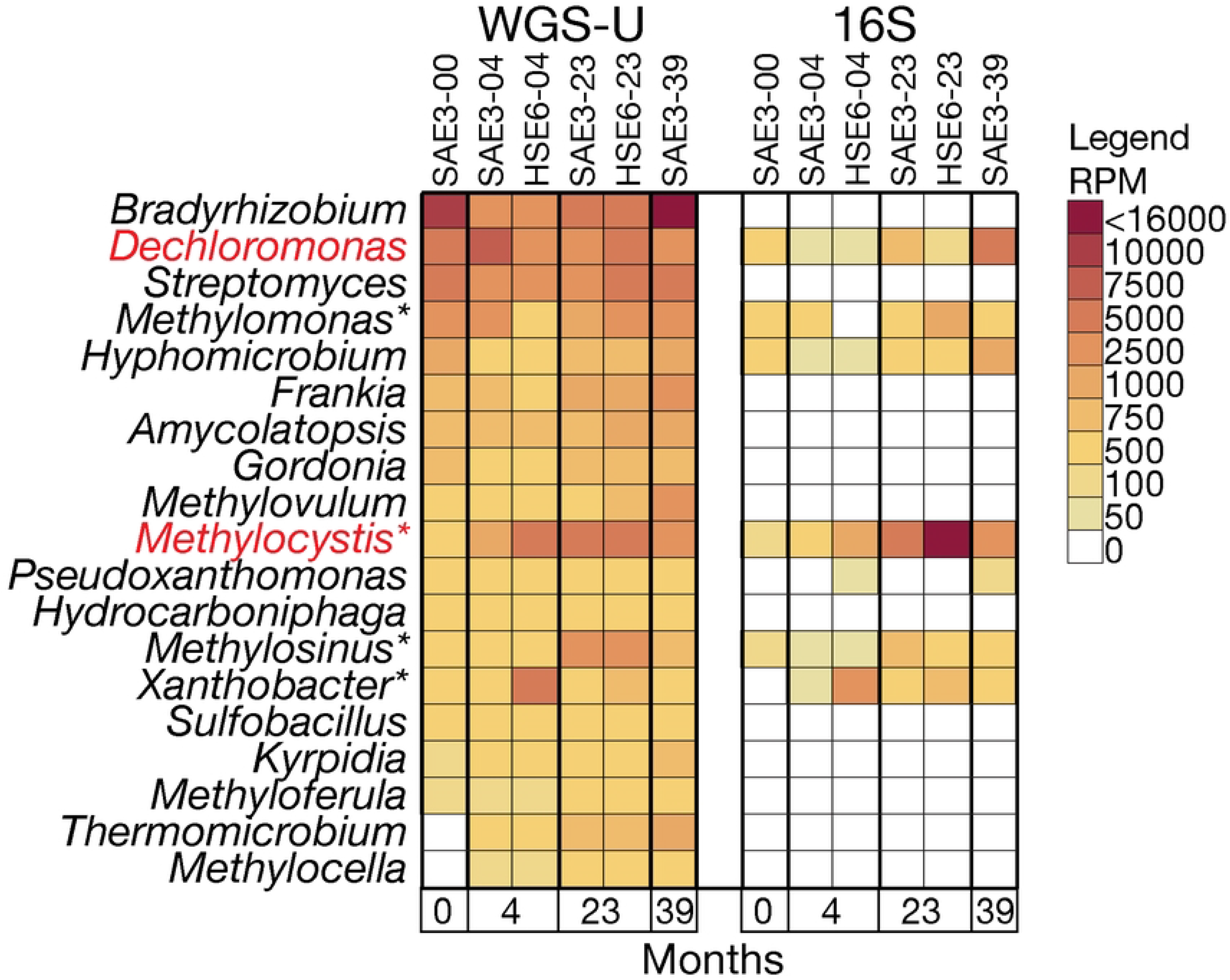
FIP-Selected Aerobic COMG. These genera include IMG/M genomes with either *mmo* or *tmo* genes that increase in abundance at NRAP.

The abundances of selected methanogenic *Archaea* are shown in Fig. 10A. *Methanoscarina* was not detected by 16S rRNA at baseline and *Methanolobus* was undetectable by 16S rRNA gene amplification. Fig. 10B are examples of genera capable of oxidating TCE due to the detection of *mo* genes at NRAP. *Xanthobacter* was undetectable at baseline for 16S rRNA gene amplicons.

**Fig. 10.**
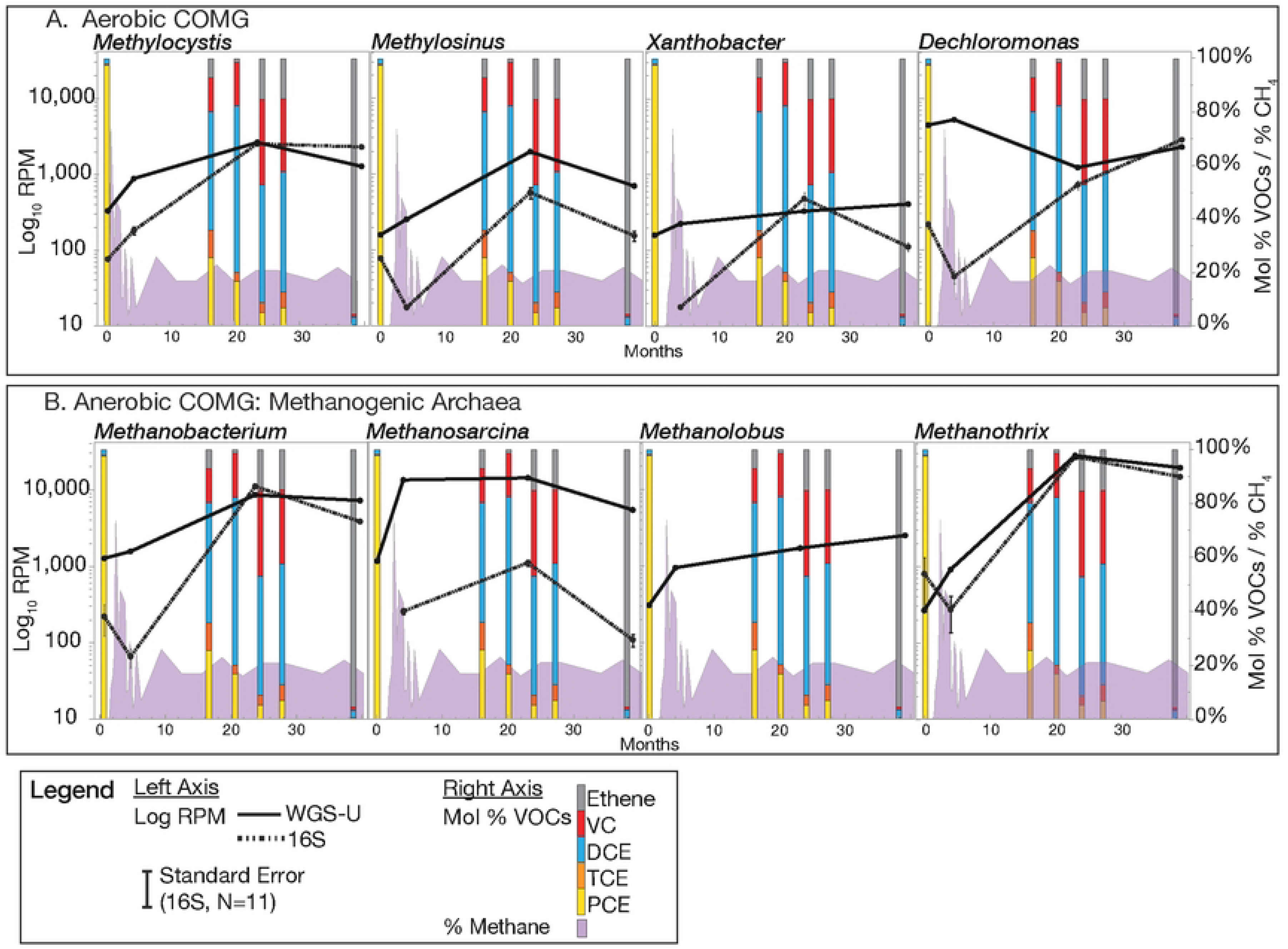
Selected COMG. Methane levels are shown since both methanotrophs and methanogens are involved. A. Methanogenic *Archaea*. B. These genomes have *mo* genes in the IMG/M database and have the potential to oxidize TCE and includes methanotrophs.

Evidence of a syntropic relationship involving taxa capable of producing O_2_ is provided in Fig. 11. These genomes contain chlorite dismutase genes in IMG/M and increase in abundance at NRAP.

**Fig. 11.**
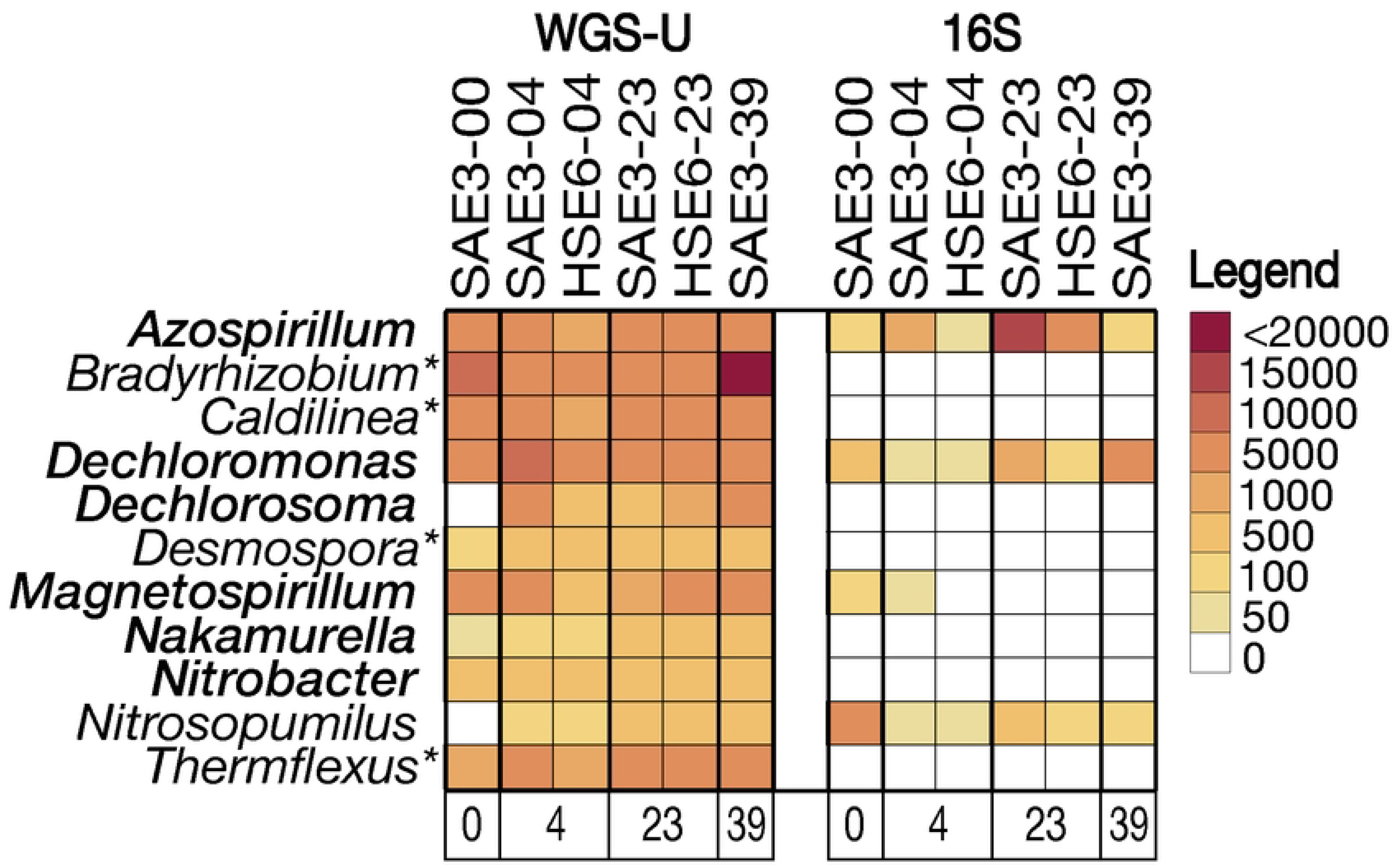
Dark O_2_ Producers at NRAP. The asterisks indicate genera known in the literature to produce dark O_2_. Those in red font have a chlorite dismutase gene-containing scaffold at NRAP.

Oxidation products and subsequent byproducts may become substrates for *hdh* gene products. Of the 212 *hdh* genes detected at NRAP, 48.5% were haloalkane, 40.7%% were haloacid, and 10.8% were haloacetate dehalogenases. These genes were distributed among 48 genera that increase at least one timepoint at NRAP and included *Geobacter*, *Trichlorobacter*, *Sulfurospirillum* and *Desulfomonile*, *Methanothrix*, *Methylocystis*, *Dechloromonas*, and *Xanthobacter*. The remainder of the *hdh*-containing genera at NRAP are shown in Fig 12.

**Fig. 12.**
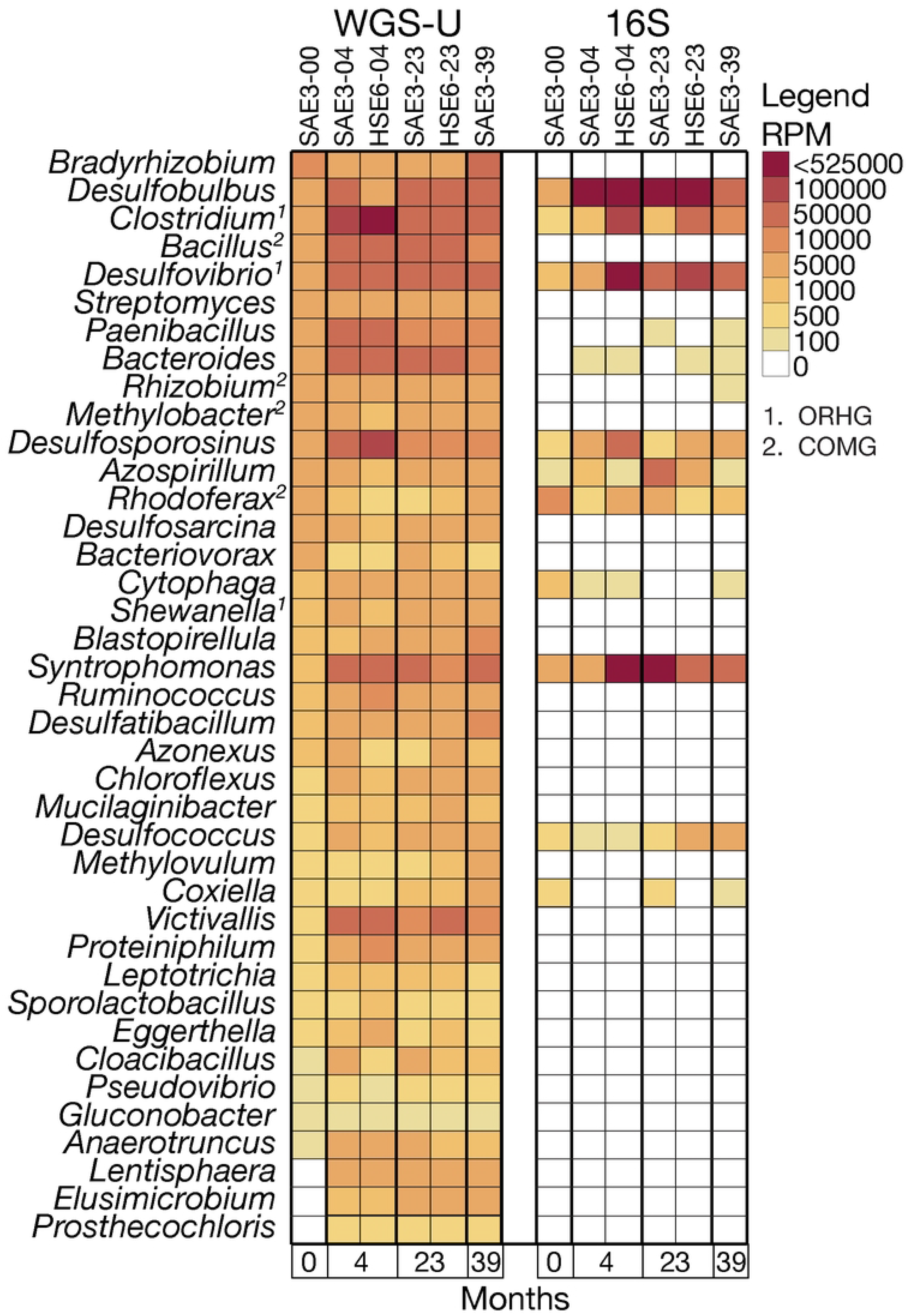
Additional Genera with *hdh*-Containing Scaffolds at NRAP. The list excludes eight previously described genera. Superscripts indicate literature-validated OHRG or COMG.

## Discussion

The L filtered on-site for each sample is negatively correlated to the biomass as measured by µg DNA/L (Fig 2A), indicating that clogging was due to an increase in biomass rather than an increase in sediment. HSE6 had the highest biomass after four and 23 months which is likely related to the initial treatment with dairy whey during the pilot phase and subsequent switch to EVO. SAE3 was exposed only to EVO.

Sampling a Superfund site has issues not encountered in laboratories, such as site access and dependence on extraction well function at sampling time. For example, HSE6 was not operational for the 39-month sampling event. Additionally, EPA regulations prevent the removal from the site the large volumes of water required to account for the low biomass environments, especially those that exist prior to remedy application. The optimization of on-site filtration methods for WGS methods is essential for field studies where transporting large volumes of water and sediment is illegal or impractical.

From the environmental engineer’s perspective the baseline is critical since it is used to determine the remedy protocol but the low biomass encountered prior to biostimulation makes it the most challenging sample to obtain. HSE6-00 did not yield enough DNA for WGS and only one 16S rRNA gene amplicon replicate was obtained (Fig. 2A). Although the same sequencing depth was achieved for all samples, there were more than 10-fold fewer WGS-U reads IDed at baseline (Fig. 2B). This finding is consistent with the expectation that WGS reveals previously undetectable diversity, known as the microbial dark matter [31]. Although all timepoints for SAE3 WGS-A had the same sequencing depth (10^10^ bp), the number of scaffolds was 100-fold lower in the baseline, but this was compensated for by the increased average scaffold size (Fig. 2C). The absence of *Dhc* and *Dhb* at baseline in WGS-A indicates that the assembly algorithm needs to be refined for a low biomass/high richness sample. Due to this discrepancy, only the combined assembly, which consolidates reads for all six samples, was further analyzed. Table 2 indicates that richness, as measured by the number of OTUs at almost all taxonomic levels and for WGS methods, is greatest at baseline. Prior to biostimulation there was lower biomass (Fig. 2A), more uncharacterized diversity (Fig. 2B), and the fraction that was IDed is the most diverse (Table 2). This is the expected response of microbiomes to ERD protocols, taxa that can adapt to anaerobic, reduced conditions have a selective advantage over other members of the diverse consortia that exist at baseline.

Both amplicon and WGS document the abundance increase of *Archaea* after remedy application that was consistent with the increase in methane on-site. (Fig. 3). WGS coverage of *Euryarchaeota* was much greater than 16S rRNA but both methods revealed increases in *Methanomicrobia* and *Methanobacteria* (Fig. 4). The baseline presence of methanogenic *Archaea* indicates the potential for anaerobic cometabolism but also predicts that mitigation procedures for the onsite build-up of methane are necessary. Validated anaerobic COMG at NRAP include *Methanobacterium*, *Methanosarcina*, and *Methanolobus* (Fig. 10A). The abundance increase is most pronounced for *Methanosarcina*, which is also capable of reverse methanogenesis under some conditions, in which CH_4_ is converted to CO_2_ [46]. Although *Methanothrix* is not yet a validated COMG, its abundance profile is consistent with that of other methanogens. The linkage between the potential of these microbes to degrade OH via transition-metal cofactors [18] and methane production remains to be determined and is critically important for balancing successful remediation with the release of methane..

Phylum-level abundance profiles fit expectations for a reduced dechlorinating environment (Fig. 4). Although *Pseudomonadota* decreases in abundance over time, both methods document increases in *Deltaproteobacteria,* which contains *Geobacter*, and *Epsilonproteobacteria,* which includes *Sulfurospirillum*. *Bacillota* class *Clostridia* (includes *Dehalobacter*) exhibits increased abundance. *Chloroflexota* abundance increases, which was expected since *Dehalococcoidia* is the only class containing OHRGs capable of complete respiration of PCE. The 2^nd^ highest abundance at baseline is *Bacteroidota*, which increases at NRAP. The presence of *Bacteroidota* is associated with improved dechlorination of TCE contaminated site undergoing bioaugmentation with KB1 [47]. *Bacteroidota* also contains halogenating enzymes, as do *Verrucomicrobiota* and *Lentisphaerota* [5], further evidence for a linkage to the biogeochemical Cl cycle.

Merging literature-validated OHRG with *rdh*-containing genomes from IMG/M indicates that only 8.2% of *rdh*-containing genome are validated chlorinated ethene reducers, reflecting the diversity of substate specificities that exist for Pfam13486 but also highlighting the need to expand OH respiration testing. Cryptic functions may also exist with amino acid profile of PFAM 13486, increasing the likelihood of misclassification. The 34.4% of literature-validated OHRG that do not have *rdh* in any reference genome for the genus may be attributable to cryptic taxa in cultures used for testing and/or cryptic pathways for OH degradation. Some taxa may harbor *rdh*-containing extrachromosomal elements that escape genome assembly. Mapping the NRAP data to the *rdh*-containing genomes of validated genera including *Dhc*, *Dhb*, *Dehalogenimonas*, *Desulfomonile* and *Trichlorobacter*. WGS-A reveals two new candidates, *Desulfobacula* and *Desulfocarbo* (Table S7).

*Dhc* exhibited a slower initial increase in WGS and decreased in 16S rRNA but both increase as VC increases (Fig 6A), consistent with results from microcosms originating from OH contaminated groundwater in which *Dhb* dominated during the presence of PCE, TCE, and DCE (also ethanes). *Dhc* increased as VC became available [9, 48]. *Dehalogenimonas* is not detected by 16S rRNA and the WGS results do not rule out a role in OH respiration at NRAP. The results for *Desulfobacula* indicate an increase from baseline to 39 months but the ability of this *Deltaproteobacteria* to catalyze OH-respiration remains to be validated.

*Geobacter* is one of the most abundant genera at NRAP, although a recommended taxonomic change moves *rdh*-containing genomes into the genus *Trichlorobacter* (Fig. 6B). *Geobacter* is known for interactions with other microbes, including the transfer of electrons along specialized pili and the production of cobalamin that can be transferred to cobalamin auxotrophs, such as *Dhc* [12, 49]. *Sulfurospirillum*, *Acetobacterium* and *Desulfitobacterium* are also cobalamin producers that increase at NRAP [13, 14]. *Desulfomonile* is a validated OHRG [50]. All eight genera in Fig.10 also harbor *hdh* genes, suggesting that these bacteria may have multiple roles in the transformation of chlorinated solvents. *Geobacter* and *Sulfurospirillum* exhibit a higher abundance in 16S rRNA amplicons than WGS-U, which is reverse of most other comparisons, evidence for amplification bias. At four months, 16S rRNA amplicon analysis results revealed large increases in the abundance of some taxa including *Geobacter* and *Sulfurospirillum*, relative to that of other taxa. WGS analysis results were proportionate. However, 16S rRNA results for other taxa including *Dechloromonas* (Fig. 10B), *Methylosinus* (Fig. 10B), *Methanothrix* (Fig. 10A), *Methanobacterium* (Fig. 10A), *Desulfomonile* (Fig 6B), *Desulfobacula* (Fig. 6A), and *Dehalococcoides* (Fig. 6A) exhibited a decrease in abundance contrary to WGS results from the four-month eDNA samples. These results suggest that high abundance taxa, such as *Geobacter* and *Sulfurospirillum,* can outcompete other taxa for the shared universal primers.

Combining known OH-degradation profiles of strains with their abundances at NRAP reveals a pathway for the complete respiration of PCE to VC (Fig. 7). The presence of *Dhc mccartyi* 195 reads at baseline is matches the original qPCR results that predicted the success of ERD at NRAP. The *Dhb* and *Dhc* scaffolds in Fig. 8 include at least one *rdhA* gene and contain accessory genes whose products are necessary for OHR [51]. *Dhc* strain 11a5 contains a potential VC reductase gene, consistent with the qPCR results and a transposase open reading frame (OrfB), suggesting that this region has the potential to be relocated to other regions in the genome, including extrachromosomal elements that can eventually be inserted into to other genomes [52]. The *MarR* transcriptional regulator is transcribed in the opposite direction from the *rdhA* genes in *Dhc* strain MB, consistent with its proposed role as a repressor of *rdh* gene transcription in *Dhc* [53]. These scaffolds are not complete gene clusters but corroborate findings that multiple *rdh* genes exist in the dechlorinating microbial consortium KB1 and from a polychlorinated biphenyl contaminated site [54, 55, 56]. Although *rdh*-containing scaffolds for *Dhc* strains 11a5 and MB are detected in the combined assembly, these strains are not identified in the WGS-U data for individual timepoints, highlighting issues of sequencing depth and differences in read identification protocols between methods.

Cometabolic pathways not linked to energy production can occur in anaerobic and aerobic environments that coexistence at OH-contaminated sites [57] and NRAP is no exception. In addition to the detection of methane and methanogenic *Archaea* at NRAP, genera with the potential for aerobic cometabolism also increase in abundance (Fig. 9). Methanotrophs *Methylocystis* and *Methylosinus* are good candidates for anaerobic cometabolic functions (Fig. 10B), but this remains circumstantial since epoxides are unstable and difficult to assay in the field. Although the increase in methane on-site can be attributed to methanogenic *Archaea*, the decrease in methane cannot be solely attributed to the action of methanotrophs since safety concerns required the installation of a vapor extraction system to collect and burn off excess methane. *Xanthobacter* and *Dechloromonas* harbor *mo* genes, but only *Xanthobacter* is a literature-validated COMG. The source of O_2_ in an anaerobic environment remains an enigma. Subsurface microenvironments, such as the soil-water interface, may sustain an O_2_ gradient or metabolic O_2_ may be available from microbial dismutation [24]. *Dechloromonas* is a facultative anaerobe known for its ability to degrade aromatic chlorinated solvents under anaerobic conditions, it can oxidize OH, and can produce O_2_ by chlorite dismutation.[58, 24]. The presence of *Dechloromonas* at NRAP is likely influenced by the presence of legacy petroleum hydrocarbon contamination in this aquifer.

The data generated for the baseline and four-month time points with WGS-U are consistent with previous NRAP results [34]. Quantification of OHRG and COMG by WGS provides an increased level of confidence since only those genomes that are validated in the literature and/or in databases as containing *rdh* genes are considered, while genus-level 16S rRNA gene amplicon results potentially include strains or genomes that do not contain *rdh* genes. The use of genome and gene-specific primers in addition to improved universal primers reduces this problem for PCR, but amplification biases remain. With adequate coverage, the scaffolding process increases confidence in the identification and quantification of OHRG and COMG by WGS when the genes for key enzymes are detected.

The use of qPCR on baseline samples to detect taxa and genes capable of OH respiration is an effective tool for remedy decisions but PCR-based techniques require an *a prior*i knowledge of microbial DNA sequences: therefore they do not facilitate the discovery of new taxa, genes, or the investigation of the metabolic pathways capable of contaminant reduction (Fig 13). WGS is an alternative to PCR that is more costly and requires more DNA but is needed to improve models to predict OH remediation. A limitation of environmental metagenomics is the lack of standardization protocols, which makes it difficult to establish parameters such as minimum detection limit [59]. The development of control samples for environmental metagenomes will enhance site-to-site comparisons. The existing DNA sequences obtained from PCR and WGS methods can be subjected to re-analysis as reference sequence databases are updated but will be informative only if metadata are available, including detailed geochemical data. The application of metabolomics to environmental microbiomes will require data from multiple sites undergoing ERD.

**Figure 13.**
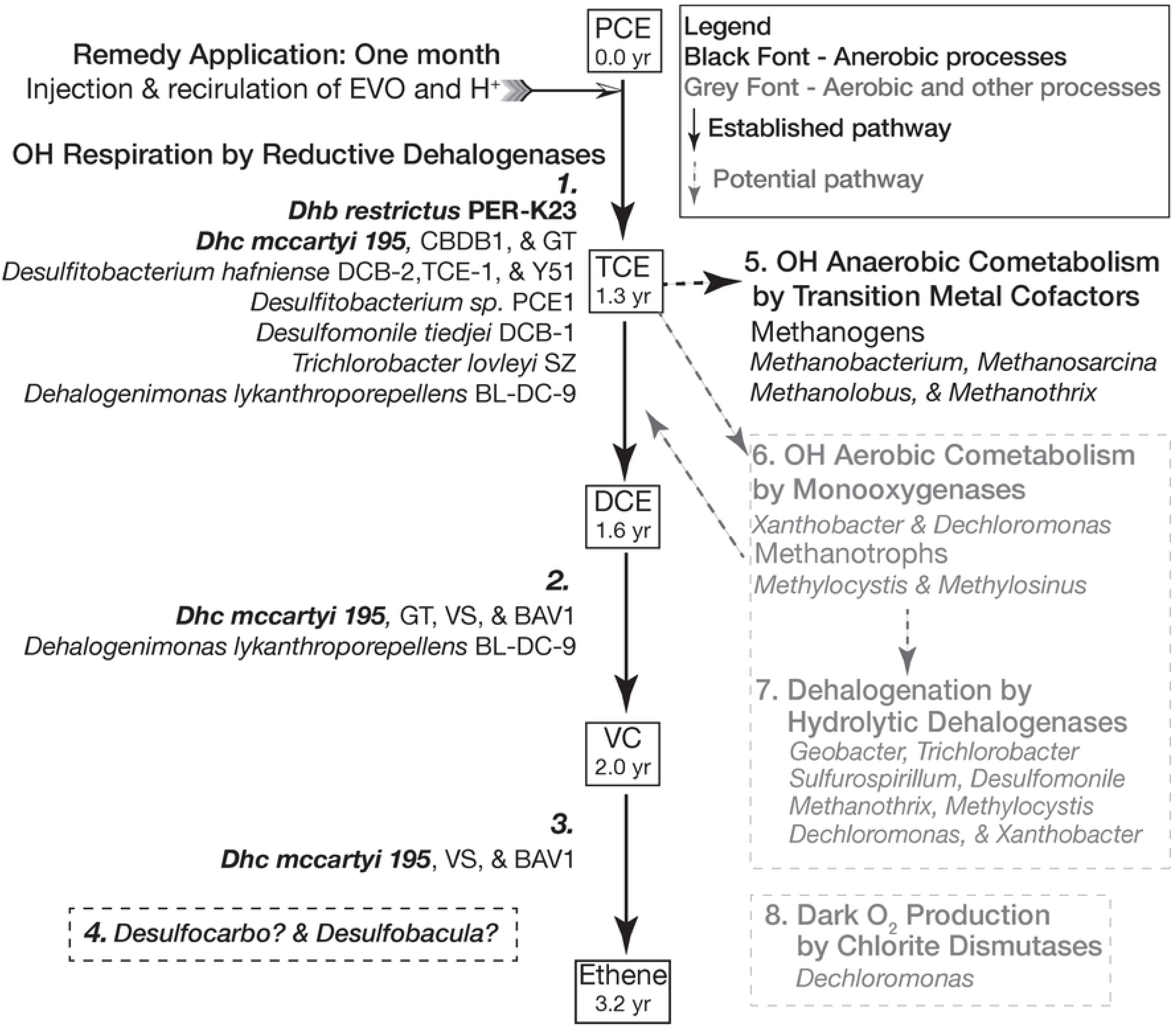
The NRAP Model for Successful ERD. The goal of ERD is the complete respiration of OHs and the presence of *Dhc*195 and *Dhb* (in bold) determined by qPCR indicated that biostimulation was the remedy of choice. The numbers shown in the pathway indicate when that compound reached its peak in the first 3.25 yrs. of the project. Relevant microbes are indicated in the number lists and their presence is supported by WGS. **1.** PCE and TCE respiring strains **2.** DCE respiring strains. **3.** VC respiring strains. **4.** Microbes that harbor *rdh* genes but are not validated as OHRG. TCE can be subjected to anaerobic or aerobic cometabolic processes. **5.** Methanogenic *Archaea*. **6.** Methanotrophs and other *mo*-containing microbes can oxidase TCE, eliminating the double bond and producing epoxides. **7.** Hydrolytic dehalogenases are abundant in the NRAP microbiome, including in microbes that play other roles in dechlorination. The potential for products produced by aerobic cometabolism, such as chlorinated ethanes, remain to be verified. **8.** The O_2_ for aerobic cometabolism may be produced by microbes that harbor chlorite dismutase.

Short read data is currently the least expensive method for WGS and its shortcomings can be mitigated by alignment into scaffolds and/or the use of long-read technology. Next on the horizon as a tool for environmental engineers is the high-throughput chromosome conformation capture (Hi-C) method (a.k.a. proximity-ligation) in which the eDNA is first cross-linked in an aliquot of the microbial community prior to cell lysis [60], [61]. A comparison of data obtained for the cross-linked sample with standard processing protocols provides a method to accurately map reads to individual genomes. Currently underway are efforts to refine on-site collection protocols to obtain a 16-year microbiome sample from the deep zone. Once sequencing is available, sequence all NRAP data will be re-analyzed using the latest reference databases and algorithms.

## Conclusions

The NRAP results demonstrate the potential of WGS to refine treatment options for contaminated groundwater and simultaneously highlight challenges associated with this technology. Among the challenges are the development of on-site sampling protocols that include guidelines relating biomass to sample volume necessary for adequate eDNA yield, the establishment of statistical procedures that include detection limit methods, read alignment algorithm refinement for low biomass/high complexity conditions, and strict data management procedures.

## Acknowledgements

Special thanks to the National Center for Genome Resources (NCGR) in Santa Fe, NM for providing preliminary sequence data.

## Supporting information

**S1 File. Tables S1 to S7 and associated references**. **Table S1**. Literature Validated OH Respiring Genera. **Table S2.** Literature-Validated OH-Cometabolizing Genera, **Table S3.** Lineage Correction Protocol, **Table S4.** Phyla Limited to Baseline WGS-U, **Table S5**. Other Candidatus Phyla in WGS-U, **Table S6.** References for OH Respiring Strains Detected at NRAP, **Table S7**. *rdh* Scaffold Parameters. Details for entries in bold are in Fig. 8. **S1 References.**

**S2 File. NRAP Data Files.** Raw and derived data. **S2.0** Read Me tab contains abbreviations and explanations for the other three tabs, **S2.1** WGS-U Master Strain, **S2.2** 16S Master Genus, **S2.3** WGS-U + 16S by Genus.

**S3 File. NRAP Figure Files.** Data for figures. **S3.0** Read Me tab contains abbreviations and explanations for the other 13 tabs; **S3.1** Biomass Fig 2A, **S3.2** WGS-U Fig 2B, **S3.3** Scaffolds Fig 2C, **S3.4** Domain & CH_4_ Pies Fig 3, **S3.5** Phylum Cell Plot Fig 4, **S3.6** Class Cell Plot Fig 4, **S3.7** FIP OHRG Fig 5, **S3.8** Selected OHRG Fig 6, **S3.9** OH Profile Strains Fig 7, **S3.10** FIP COMG Fig 9, **S3.11** Selected COMG Fig 10, **S3.12** Dark O_2_ Fig. 11, **S3.13** FIP *dh* Fig 12.

**S4 File. NRAP Genes and Genomes.** Database-derived and other data. The **S4.0** Read Me tab contains abbreviations and explanations of the other nine tabs, which are as follows: **S4.1** MI Results, **S4.2** Lit Val OHRG & COMG, **S4.3** IMG *rdh* genomes, **S4.4** IMG *mo* genomes, **S4.5** NRAP IMG *rdh* genus, **S4.6** IMG NRAP *mo* genus, **S4.7** NRAP *rdh* scaffolds, **S4.8** NRAP *dh* scaffolds, **S4.9** NRAP *hdh* by genus

